# Homozygous NMNAT2 mutation in sisters with polyneuropathy and erythromelalgia

**DOI:** 10.1101/610907

**Authors:** Peter Huppke, Eike Wegener, Jonathan Gilley, Carlo Angeletti, Ingo Kurth, Joost P.H. Drenth, Christine Stadelmann, Alonso Barrantes-Freer, Wolfgang Brück, Holger Thiele, Peter Nürnberg, Jutta Gärtner, Giuseppe Orsomando, Michael P. Coleman

**Affiliations:** Department of Pediatrics and Pediatric Neurology, University Medical Center Göttingen, Georg August University Göttingen, Germany.; John van Geest Centre for Brain Repair, University of Cambridge, ED Adrian Building, Forvie Site, Robinson Way, Cambridge, CB2 0PY, UK.; Babraham Institute, Babraham Research Campus, Babraham, Cambridge CB22 3AT, UK.; Department of Clinical Sciences (DISCO), Section of Biochemistry, Polytechnic University of Marche, Via Ranieri 67, 60131 Ancona, Italy, Email; Institut für Humangenetik, Universitätsklinikum Jena, Kollegiengasse 10, 07743 Jena, Germany; Department of Gastroenterology & Hepatology, Radboud UMC, P.O. Box 9101, 6500 HB Nijmegen, Netherlands; Institute of Neuropathology, University Medical Center, Georg August University Göttingen, Germany; Department of Neuropathology, University Medical Center Leipzig, Leipzig, Germany; Cologne Center for Genomics (CCG), University of Cologne, Cologne, Germany; Center for Molecular Medicine Cologne (CMMC), University of Cologne, 50931 Cologne, Germany

## Abstract

We identified a homozygous missense mutation in the gene encoding NAD synthesizing enzyme NMNAT2 in two siblings with childhood onset polyneuropathy with erythromelalgia. No additional homozygotes for this rare allele, which leads to amino acid substitution T94M, were present among the unaffected relatives tested or in the 60,000 exomes of the ExAC database. For axons to survive, axonal NMNAT2 activity has to be maintained above a threshold level but the T94M mutation confers a partial loss of function both in the ability of NMNAT2 to support axon survival and in its enzymatic properties. Electrophysiological tests and histological analysis of sural nerve biopsies in the patients were consistent with loss of distal sensory and motor axons. Thus, it is likely that NMNAT2 mutation causes this pain and axon loss phenotype making this the first disorder associated with mutation of a key regulator of Wallerian-like axon degeneration in humans. This supports indications from numerous animal studies that the Wallerian degeneration pathway is important in human disease and raises important questions about which other human phenotypes could be linked to this gene.

## INTRODUCTION

Erythromelalgia is a rare clinical syndrome characterized by intermittent attacks of intense burning pain with redness and swelling predominantly affecting the lower extremities but which may also be present in upper extremities. It was named by S. Weir Mitchell in 1878 from erythro (red), melos (extremity), and algos (pain)^1^. The episodes can be induced by heat, exercise and gravity and can be relieved by cooling and elevation. They usually have their onset in the first decade but may also first appear during adulthood. Erythromelalgia can be classified as primary or secondary. In a population-based study from Minnesota, the incidence of erythromelalgia overall was 1.3 per 100,000 people per year with a preponderance of primary erythromelalgia (1.1 per 100,000 people per year)^2^. Secondary erythromelalgia can occur in a number of disorders like small fiber peripheral neuropathies, thrombocythemia, myeloproliferative diseases, as a paraneoplastic syndrome or as a side effect of different medications^3^. Many patients affected with primary erythromelalgia carry gain-of-function mutations in the *SCN9A* gene coding for the Nav1.7 sodium channel^4^, either transmitted in an autosomal dominant manner or as de novo mutations. Nav1.7 is expressed in small-diameter nociceptive neurons where it amplifies small depolarizations bringing the membrane potential closer to the voltage threshold of an action potential^5^.

We diagnosed small fiber neuropathy in two sisters suffering from childhood onset polyneuropathy with erythromelalgia. Sequencing of the complete coding region of *SCN9A* did not reveal any mutations. We therefore proceeded to exome sequencing in both sisters and the parents and discovered a homozygous mutation in *NMNAT2* in the patients and heterozygosity in the parents. Biochemical and cell biological analyses indicate a partial loss of NMNAT2 function caused by the mutation, consistent with the hypothesis that these mutations are causative for the phenotype seen in the siblings.

## MATERIALS AND METHODS

### Exome Sequencing, Filtering and bioinformatic prediction

DNA samples were obtained from patients and parents following informed consent and approval by local ethic committee from the University Medical Center Göttingen, Göttingen, Germany. DNA was extracted from peripheral EDTA blood using standard protocols. For each exome, 1 µg of DNA was fragmented, barcoded and enriched using the NimbleGen SeqCap EZ Human Exome Library v2.0 enrichment kit (Roche NimbleGen, Madison, WI, USA). Purified and quantified library pool was subsequently sequenced on an Illumina HiSeq 2000 sequencing instrument (Illumina, San Diego, CA, USA) using a multiplex paired end 2×100bp protocol. Data processing, analysis and filtering were performed using the ‘Varbank’ GUI and pipeline version 2.14 (CCG, University of Cologne, Germany) (https://varbank.ccg.uni-koeln.de/). Reads were mapped to the human genome reference build hg19 using the BWA-aln alignment algorithm, resulting in ≥30× coverage for 88.5% and 89.7% of target sequences in the two exomes, respectively. GATK v.1.62 was used to mark duplicated reads, to do a local realignment around short insertions and deletions, to recalibrate the base quality scores and to call SNPs and short Indels. The GATK UnifiedGenotyper variation calls were filtered for high-quality (DP>15; AF>0.25; QD>2; MQ>40; FS<60; MQRankSum>-12.5; ReadPosRankSum>-8; HaplotypeScore <13) rare (MAF ≤ 0.005 based on 1000 genomes build 20110521 and EVS build ESP6500 and the Exome Aggregation Consortium (http://exac.broadinstitute.org/)) variants, predicted to modify a protein sequence or to impair splicing, implicated by reduced maximum entropy scores (MaxEntScan). False positive and irrelevant variants were further reduced by taking advantage of the Varbank InHouseDB containing 511 epilepsy exomes. Compound heterozygous and homozygous/hemizygous variants were extracted for the patient. Prediction of functional impact of all received variants was performed using the dbNSFP version 3.0a36,37. DbNSFP software co-applied several in silico analysis tools to predict the conservation, using PhastCons and GERP, and the functional consequence, using SIFT, PolyPhen2, Provean, LRT, MutationTaster, MutationAssessor, FATHMM, VEST, MetaSVM and MetaLR, of the affected site. In addition phenotype genotype correlations were investigated using public database Online Mendelian Inheritance (OMIM) (http://www.omim.org/), Orphanet (http://www.orpha.net) and ClinVar (https://www.ncbi.nlm.nih.gov/clinvar).

Mutations found by NGS were confirmed by PCR and Sanger-Sequencing. For PCR, genomic DNA and the following primers used: Forward 5’-TCT AAG CGC ATG GTA TGG C3’ and Reverse 5’-CTT TCA CTC CCT CCT CCT TTG-3’. Gel electrophoresis was performed using a 1% agarose gel. For sequencing, specific bands were eluted from the gel by use of NucleoSpin^®^ Gel and PCR Clean-up (Machery-Nagel, Düren, Germany) and subsequently processed for direct dye terminator sequencing with BigDye Terminator Ready Reaction chemistry 3.1 on an ABI PRISM 3130-Avant genetic analyser (Life Technologies, Darmstadt, Germany). All reactions were performed according to instructions provided by the manufacture.

### Constructs

NMNAT2 T94M and H24D mutations were introduced by QuikChangeII site-directed mutagenesis (Stratagene) into the complete open reading frame (ORF) of the canonical 307 amino acid human NMNAT2 variant (NCBI ProteinID: NP_055854) cloned in expression vector pCMV Tag-2B (Stratagene). A short linker (17 amino acids) is present between and N-terminal tag Flag and the NMNAT2 ORF. DNA sequencing (Cogenics) confirmed the presence of the T94M mutation and the absence of PCR errors. The wild-type or T94M mutant ORFs were then PCR-cloned into pET28c (Novagen) for bacterial production of the proteins with an N-terminal 6× His tag. pDsRed2-N1 (Clontech) was used for expression of variant *Discosoma* red fluorescent protein (DsRed2) to fluorescently-label injected neurons and their neurites. A SARM1-GFP expression vector consisting of the entire coding region of murine SARM1 cloned (by PCR) into pEFGP-N1 (Clontech, C-terminal eGFP tag) was used for re-expression of SARM1 in *Nmnat2*^*gtE/gtE*^;*Sarm1*^−/−^ neurons. pEGFP-C1 (Clontech) was used to express enhanced green fluorescent protein (eGFP) in HEK 293T cells to act as a transfection control / reference for NMNAT2 turnover.

### Microinjection, transection of neurites and imaging

Dissociated SCG neuron cultures (from P0-P2 mouse pups), expression vector injections, Flag immunostaining and the quantification of neurite survival after transection were all performed essentially as described previously^6,7^. Expression vectors and the concentrations used in injection experiments are described in the Figure 5 legend. Fluorescence images were acquired as described previously^6^. For transection assays at 39°C, cells were cultured at 37°C up to the time of transection and were only then transferred to 39°C for the duration of the assay (24 hours). Mean intensities of Flag immunostaining and DsRed fluorescence signals in injected SCG neurons were determined from images captured with a 4× objective using Fiji software http://fiji.sc^8^ by thresholding (30 and 18 respectively, dark background) followed by particle analysis to identify neurons with signal intensity above background (threshold value was subtracted from mean intensity values obtained).

### HEK 293T transfection

HEK 293T cells were cultured in DMEM with 4,500 mg/L glucose and 110 mg/L sodium pyruvate (PAA), supplemented with 2 mM glutamine and 1% penicillin/streptomycin (both Invitrogen), and 10% fetal bovine serum. Cells were plated in 24-well format to reach 50-60% confluence before transfection with Lipofectamine 2000 reagent (Invitrogen) according to the manufacturer’s instructions. 500 ng Flag-NMNAT2 expression vector (wild-type or T94M), 200 ng of an empty pCMV-Tag series vector, and 100 ng pEGFP-C1 were transfected per well. 10 µM emetine hydrochloride (Sigma-Aldrich) was used to block protein synthesis. Cells from single wells at each timepoint after treatment were lysed directly in 100 µl 2× Laemmli sample buffer, sonicated to fragment genomic DNA in the absence of heating, and equal amounts (10µl) used for immunoblotting.

### Immunoblotting

Extracts were resolved on 4-15 or 4-20% SDS polyacrylamide gels, transferred to Immobilon-P membrane (Millipore) and probed with antibodies essentially as described previously^7^. The following primary antibodies were used: mouse monoclonal anti-FLAG M2 (1:2,000 Sigma-Aldrich F3165), mouse monoclonal anti-GFP clones 7.1 and 13.1 (1:2,000, Sigma-Aldrich 11814460001) and rabbit polyclonal α-Tubulin (1:7,500, Thermo Fisher Scientific PA5-29444). Appropriate HRP-conjugated secondary antibodies were used for band detection with SuperSignal™ West Dura Extended Duration Substrate (Thermo Fisher Scientific) using an Alliance chemiluminescence imaging system (UVITEC Cambridge). Relative band intensities on captured digital images were determined from the area under histogram peaks using Fiji software http://fiji.sc^8^.

### Bacterial expression, and purification

The above pET28c plasmid constructs of human NMNAT2 wild-type^9^ and mutant T94M encoded recombinant proteins with the same N-terminal tag: MGSSHHHHHHSSGLVPRGSH. Expression was carried out into *E. coli* BL21(D3) cells (Invitrogen) following 0.5 mM IPTG induction for 17h at 28 °C and purification by TALON chromatography (Clontech), essentially as described^10^. The purified proteins were finally desalted on PD-10 columns (GE Healthcare) in 50 mM HEPES/NaOH buffer, pH 7.5, 1 mM Tris(2-carboxyethyl)phosphine (TCEP), 20 % glycerol, and stored at −80 °C. Their concentration was measured by the Bio-Rad protein assay. Their purity was evaluated on 12% SDS polyacrylamide gels either by Coomassie staining or after immunoblotting as described above, using monoclonal anti-NMNAT2 (1:1,000 Abcam Ab5698) as the primary antibodies.

### Activity assays

One unit (U) of NMNAT activity is defined as the enzyme amount catalyzing 1 μmol/min product formation at the indicated temperature. Rates were routinely measured at 37 °C by a spectrophotometric coupled assay^11^, unless otherwise indicated, in 0.5 mL mixtures containing 30 mM HEPES/NaOH buffer, pH 7.5, 0.5 mg/mL bovine serum albumin (BSA Sigma-Aldrich A7906), 75 mM ethanol, 30 mM semicarbazide (Sigma-Aldrich S2201), 12.5 U/mL alcohol dehydrogenase (ADH Sigma-Aldrich A7011). The wild type NMNAT2 was added at 0.2-0.8 μg/mL in the presence of 25 mM MgCl_2_ and 1 mM both ATP and NMN. The T94M mutant was added at 0.5-1.5 μg/mL in the presence of 5 mM MgCl_2_, 2 mM ATP, and 1 mM NMN. The Mg^2+^-dependence was evaluated using 1-100 mM MgCl_2_ variable concentrations. The *K*_m_ and *K*_cat_ values were calculated as described^12^ using 60-800 μM ATP and 40-600 μM NMN for the mutant, or 50-600 μM and 10-150 μM respectively for the wild type. Due to the known instability of NMNAT2 preparations after thawing^10^, enzyme addition was always used to start the reaction, and control assays were performed in parallel. Further assays of both enzymes coupled to pure NAMPT (*Mus musculus* nicotinamide phosphoribosyltransferase obtained as described previously^13^ were carried out at 30 °C by HPLC. Assay mixtures (0.5 ml) contained ~7 mU/ml human NMNAT2 (either wild type or T94M), ~2 mU/ml murine NAMPT, ~100 mU/ml inorganic pyrophosphatase (PPase Sigma-Aldrich I1643) in 100 mM HEPES/NaOH buffer pH 7.5, 0.5 mg/ml BSA, 1 mM TCEP, 1mM Nam, 3 mM 5-phosphoribosyl 1-pyrophosphate (PRPP Sigma-Aldrich P8296), 2 mM ATP, and 5 mM MgCl_2_. Reactions started by adding Nam and were stopped at the indicated times by HClO_4_/K_2_CO_3_ treatment, followed by ion-pair reverse-phase C18-HPLC analysis^14^ for the quantification of Nam, NMN, and NAD.

### Statistics

Statistical testing of data was performed using Excel (Microsoft) or Prism (GraphPad Software Inc., La Jolla, USA). The appropriate tests used are described in the Figure legends. A *p* value < 0.05 was considered significant.

## RESULTS

### Polyneuropathy with erythromelalgia in two siblings

Family history: Patients 1 and 2 are the second and third children of consanguine parents. The parents, first cousins, as well as an older sister (age 21 years) are healthy. Nerve conduction studies in the mother were normal. In the extended family in Turkey no further members with erythromelalgia are known.

Patient 1 was born after an uneventful pregnancy (birthweight 2700gr, length 50cm). Development was normal (sitting 6 months, walking 11 months, first words 12 months). At age 4 years she experienced her first attack during an infection with very severe pain in both feet necessitating hospitalization for 6 weeks. The pain had a burning character that was increased by touch and only relieved by cooling. There was mild swelling and reddening of the feet (Fig. 1). Following this episode, she had recurrent similar attacks lasting up to one month. At age 10 and 12 years she again experienced very severe attacks necessitating hospitalization for several weeks. After puberty the attacks were milder and less frequent (every 2-3 months). The attacks were provoked by warm shoes, exercise or psychological stress but immersion of the feet in cold water could relieve the pain for 15-30 minutes. At age 19 she also reported stiffness in the feet and the hands, especially in the morning. A therapeutic trial with colchicine had no effect, but steroids seemed to shorten the attacks. In addition, she presented with bilateral pes cavus and pes equinus and a mild tremor of both hands (Fig. 1). Other than a reduced sensation for temperature in the feet and mild calf atrophy, the neurological examination between the attacks was normal.

**Figure 1:**
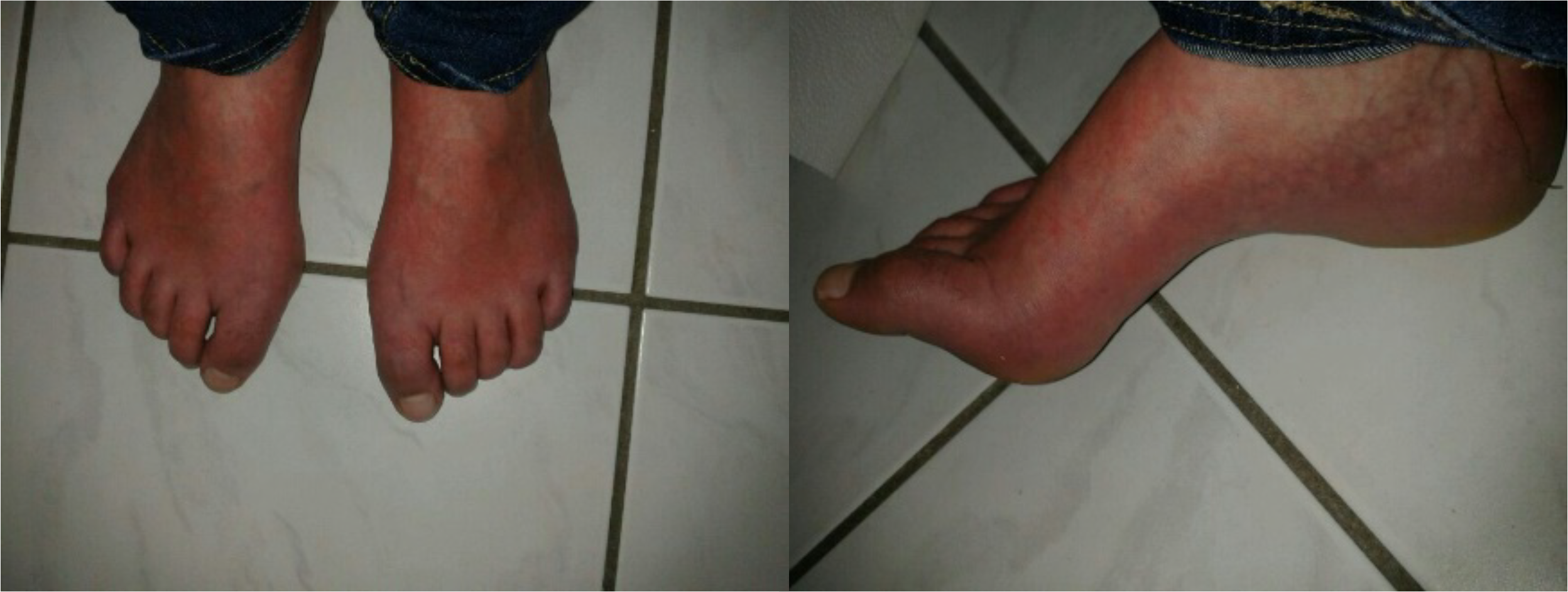
Clinical features. Pes cavus and reddening as well as swelling of the feet of patient 1 during a painful attack.

Extensive investigations in blood, CSF and urine during the attacks and in the asymptomatic intervals were normal without signs of rheumatologic or other inflammatory disorders. Normal α-galactosidase A activity ruled out Fabry disease and cranial and spinal MRI performed at age 19 were normal. Nerve conduction studies showed markedly reduced muscle action potential amplitudes for peroneal and tibial nerve and reduced sensory amplitudes for the sural and median nerve with conduction velocities being normal in both cases (data not shown), suggesting axonal motor and sensory neuropathy more pronounced in the legs than the arms. Staining of semithin transverse sections of a sural nerve biopsy taken at age 14 showed a homogeneous and moderate loss of myelinated axons with thinly myelinated fibers also seen (Fig. 2A), indicating remyelination or axonal regeneration. Ultrastructurally, a slight reduction of non-myelinated fibers was apparent and was accompanied by an increase in the number of collagen pockets (Fig. 2B). In the teased fiber preparation, myelin ovoids, suggestive of Wallerian degeneration, were found in a single fiber (Fig. 2C). Neither an increase in the number of macrophages (KiM1P) nor T lymphocytes (CD3) could be demonstrated using immunohistochemical methods (Figs. 2D and E).

**Figure 2:**
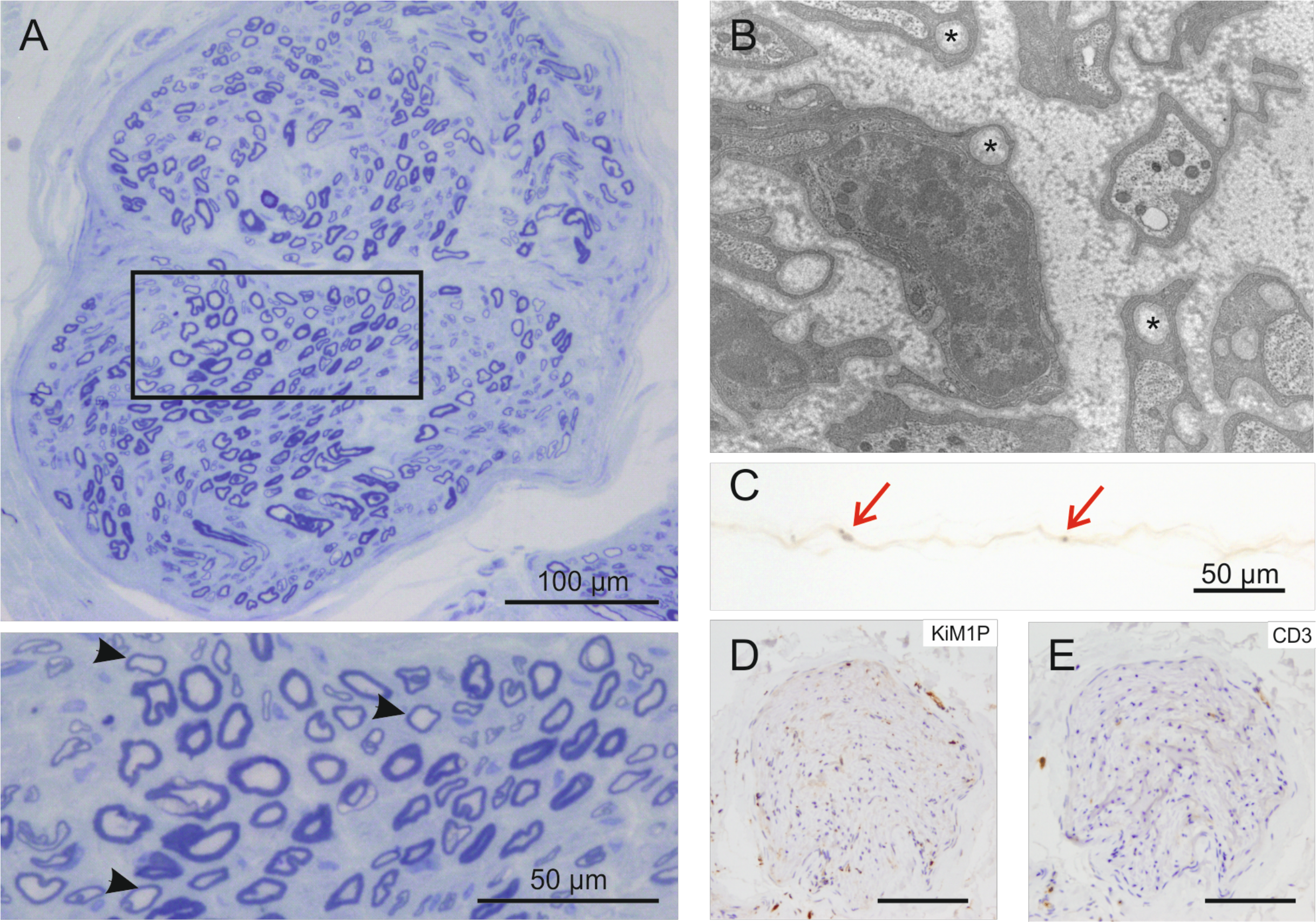
Sural nerve biopsy. A. Semithin section showing a homogeneous, moderate reduction in myelinated axons and some fibers with thin myelin sheaths, most likely reflecting remyelination or axon regeneration (arrowheads, inset). B. Electron microscopy reveals the presence of collagen pockets (asterisks). C. Teased fiber preparation showing evidence of myelin ovoids in a single fiber, indicating Wallerian degeneration (arrows). No increase in endoneural macrophages (D, KiM1P) or T lymphocytes (E, CD3), was observed. Unless stated otherwise, scale bars correspond to 100um.

Patient 2 was born 3 years after patient 1 after an uneventful pregnancy (weight 3260 g, length 48 cm). Similar to her older sister, she had normal psychomotor development. Her first episode of severe pain in the feet with mild swelling was precipitated by an infection at age 4 and lasted 4 weeks. Since then she has attacks lasting 1-7 days every 2-3 months. These episodes could be provoked by exercise, heat or infections. Overall she was less affected than her sister and with time the episodes became less severe. Physical examination at age 16 showed pes cavus and pes equinus. Electrophysiology showed axonal neuropathy with reduced muscle action potential amplitudes and normal nerve conduction velocity for motor and sensory nerves. Cranial and spinal MRI scans performed at age 16 were normal.

### A homozygous missense mutation in NMNAT2 cosegrates with the phenotype

After genetic analysis of the *SCN9A* gene did not reveal any abnormalities, exome sequencing was performed. Nonsynonymous, homozygous variants in 4 genes, TGFB1I1 (c.454C>T, p.R152C), *AHSP* (c.224A>T, p.N75I, rs75782426), LAMC1 (c.923A>G, p.K308R, rs139092535) and *NMNAT2* (c.281C>T, p.T94M) were detected in both affected sisters. The variants in the *AHSP* and *LAMC1* genes have been described previously with an allele frequency of 0.0047 (including 5 homozygous individuals) and 0.00051 (including 1 homozygous individual), respectively (gnomAD v2.1.1 [controls]) and were therefore considered unlikely to be causative for the disorder seen in the patients. The *TGFB1I1* gene codes for the transforming growth factor beta1 induced transcript 1 also known as hydrogen peroxide-inducible clone-5 (Hic-5) which is highly expressed in vascular smooth muscle cells of different organs and participates in the transcriptional regulation of several genes^15^. Mice lacking Hic-5 show no obvious abnormalities^16^. Sanger sequencing of the *TGFB1I1* gene in the index family showed that both affected sisters but also the unaffected sister were homozygous for the variant, while the parents are heterozygous (data not shown). The fourth homozygous altered gene in the two affected sisters was *NMNAT2*, which codes for nicotinamide mononucleotide adenylyltransferase 2 (NMNAT2), the predominant axonal isoform of a family of compartmentalized NAD synthesizing enzymes^17^. As the patients had signs of polyneuropathy this gene was a good candidate because NMNAT2 has been shown to be critical for axon survival in primary neuronal cultures^7^ and mouse embryos lacking NMNAT2 expression die around birth most likely due to severe truncation of peripheral nerve axons and CNS axon tracts^6,18^. Sanger sequencing of the *NMNAT2* gene in the index family showed that both parents and the unaffected sister were heterozygous for the c.281C>T variant while the two affected sisters were homozygous (Fig 3). We established a cohort of 87 patients suffering from erythromelalgia coming from Germany or the Netherlands. DNA was obtained from these patients and Sanger sequenced for SCN9A mutations. None of the patients from this cohort carried disease associated variants.

**Figure 3:**
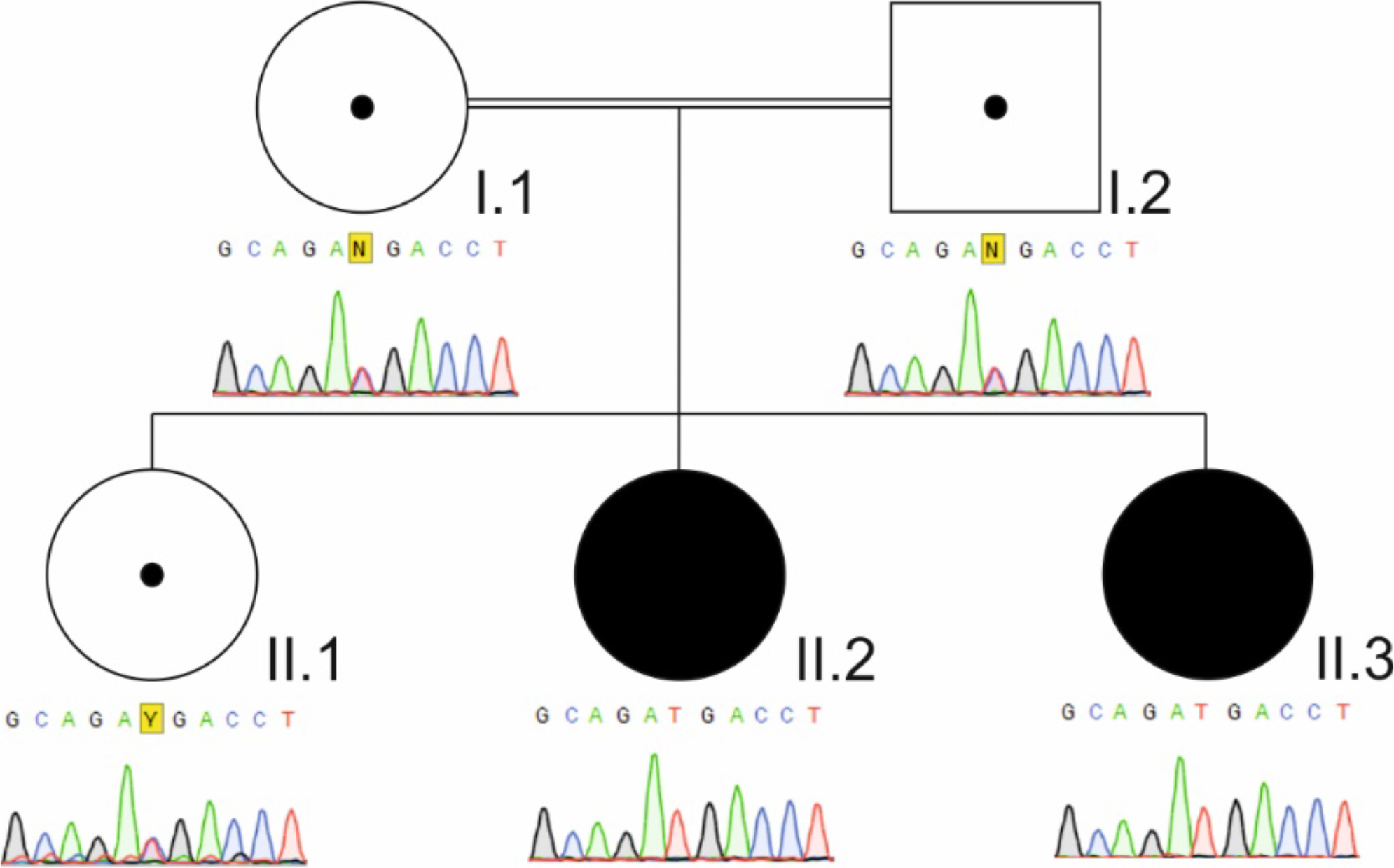
Sanger sequencing of NMNAT2 across the T94M codon. Sequencing shows heterozygosity for the c.281C>T variant in the mother (I.1) the father (I.2) and the unaffected sister (II.1) and homozygosity for the 2 affected sisters (II.3, patient 2 and II.4, patient 1).

### T94M NMNAT2 has reduced capacity to promote neurite survival that is partly temperature-dependent

To test whether the homozygous *NMNAT2* mutation in these siblings could be the underlying cause of their symptoms, we investigated whether the encoded T94M substitution might alter the ability of human NMNAT2 to promote axon survival. To do this, we assessed the relative capacity of Flag-tagged T94M NMNAT2 to delay the degeneration of transected neurites (Wallerian degeneration) in cultured mouse superior cervical ganglion (SCG) neurons using an established microinjection method that can differentiate the axon-protective competence of NMNAT proteins, including a variety of artificial NMNAT2 mutants^7,19^.

Given that pain episodes in the affected patients are often precipitated by localized or systemic increases in temperature, we first compared the ability of T94M Flag-NMNAT2 to preserve the integrity of cut neurites at both 37°C and 39°C relative to wild-type Flag-NMNAT2 and an artificial, enzyme-dead mutant, H24D Flag-NMNAT2^20^. Expression vectors for each Flag-NMNAT2 protein were introduced into wild-type SCG neurons at a relatively low concentration, at which wild-type Flag-NMNAT2 preserves ~70% of transected neurites of the injected neurons for at least 24 hours at 37°C, to provide a stringent test of their protective capacities (Fig. 4A and 4B). At this vector concentration, T94M Flag-NMNAT2 was able to protect cut neurites almost as well as wild-type Flag-NMNAT2 at 37°C but, intriguingly, there was a proportionately much greater reduction in its relative capacity to protect at 39°C (Fig. 4A and 4B). In contrast, H24D Flag-NMNAT2 conferred no protection at either temperature, as expected for an NMNAT2 protein lacking enzymatic activity^19,20^. Crucially, while expression level can strongly influence protective capacity in this type of experiment, Flag immunostaining revealed that T94M Flag-NMNAT2 expression broadly matches that of wild-type Flag-NMNAT2 in injected neurons (Fig. 4C).

**Figure 4.**
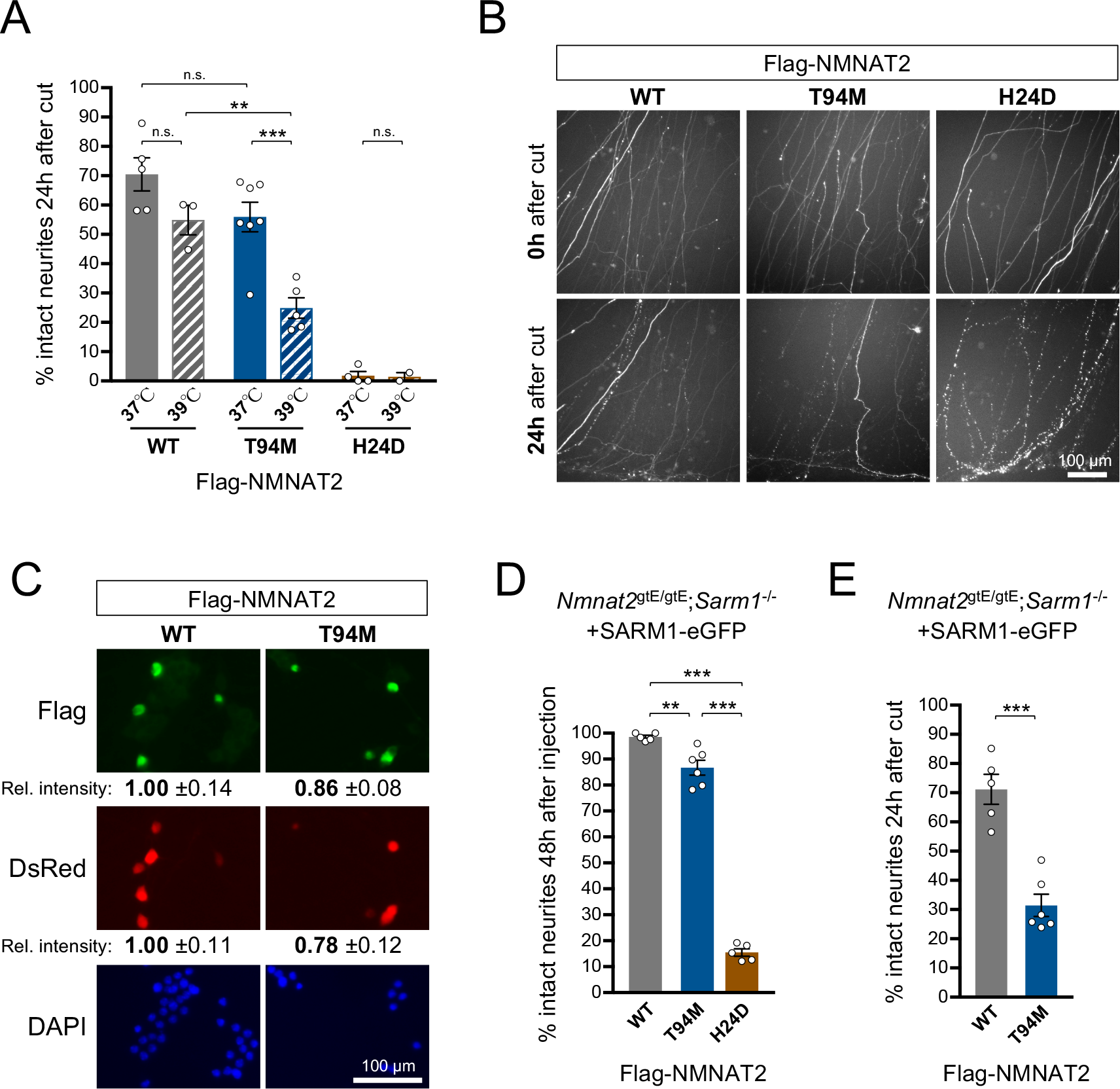
A reduced capacity of T94M NMNAT2 to maintain neurite health is partly temperature-dependent. (A) 24 hour survival of transected neurites of wild-type SCG neurons co-injected with expression vectors for wild-type (WT), T94M or H24D Flag-NMNAT2 (10 ng/µl) and DsRed (50 ng/µl) at either 37°C or 39°C. Neurites were cut 48 hours after injection (when DsRed expression is sufficient to clearly label distal neurites). Surviving, non-degenerated neurites (continuous DsRed labeling) is shown as a percentage of the number of healthy neurites at the time of cut (0h). Individual values and means ± SEM are plotted (individual values represent the average of two fields from the same culture). n.s. = not significant (p > 0.05), ** p < 0.01, *** p < 0.001, two-way ANOVA with Tukey’s multiple comparisons test. (B) Representative images of assays performed at 39°C as described in A. Images show transected neurites at 0 and 24 hours after cut with the lesion site located just below the bottom edge of the field. Brightness and contrast have been adjusted to optimize neurite visualization. (C) Relative expression of wild-type (WT) or T94M Flag-NMNAT2 in injected SCG neuron cell bodies. Representative fluorescent images are shown of SCG neurons 48 hours after co-injection with expression vectors for WT or T94M Flag-NMNAT2 (10 ng/µl) and DsRed (50 ng/µl); DsRed identifies injected neurons, Flag immunostaining shows expression of the Flag-NMNAT2 proteins, DAPI labels all nuclei. Relative intensities (± SEM) of Flag immunostaining and DsRed signal are shown after transformation to the mean of levels in neurons injected with the wild-type Flag-NMNAT2 construct. WT and T94M data was calculated from 41 and 43 injected neurons (DsRed positive) of which 65.9% and 62.8% were Flag-positive respectively. (D) 48 hour survival of (uninjured) neurites after co-injection of *Nmnat2*^*gtE/gtE*^;*Sarm1*^−/−^ SCG neurons with expression vectors for wild-type (WT), T94M or H24D Flag-NMNAT2 (10 ng/µl), SARM1 with a C-terminal eGFP tag (7.5 ng/µl) and DsRed (40 ng/µl). Surviving, non-degenerated neurites (continuous DsRed labeling) is shown as a percentage of the total number of DsRed-labeled neurites (non-degenerated plus degenerated neurites with discontinuous DsRed label and/or DsRed-positive swellings) present at 48 hours. Data is expressed as in A. ** p < 0.01, *** p < 0.001, one-way ANOVA with Tukey’s multiple comparisons test. (E) 24 hour survival of transected neurites of *Nmnat2*^*gtE/gtE*^;*Sarm1*^−/−^ SCG neurons injected as in D (the few surviving neurites of neurons injected with the H24D Flag-NMNAT2 construct were not assayed). The assay was performed at 37°C and data presented as described in A. *** p < 0.001, Student’s *t*-test.

The above experiments were performed in the presence of endogenous, murine NMNAT2, but the affected siblings, homozygous for the *NMNAT2* mutation, should only express T94M NMNAT2. Therefore, we next assessed whether the presence of endogenous NMNAT2 might mask a greater difference in protective capacity of wild-type and T94M Flag-NMNAT2 by performing the same neurite transection assay (at 37°C) in NMNAT2-deficient neurons. Neurons lacking NMNAT2 alone (*Nmnat2*^*gtE/gtE*^ neurons) are unsuitable for this due to their severely stunted neurites^6^, but NMNAT2-deficient neurons additionally lacking the pro-degenerative protein SARM1 (*Nmnat2*^*gtE/gtE*^;*Sarm1*^−/−^ neurons) grow normally and the inherent resistance of their neurites to degeneration can be overcome by SARM1 re-expression^21^. NMNAT2 enzyme activity is required for survival of intact neurites when SARM1 is present^21^ and, consistent with this, we found that the majority of (uninjured) neurites of *Nmnat2*^*gtE/gtE*^;*Sarm1*^−/−^ neurons degenerated after co-injection with expression vectors for enzyme-dead H24D Flag-NMNAT2 and SARM1(-GFP) (Fig. 4D). In contrast, (uninjured) neurites were largely preserved up to 48 hours after co-injection with T94M Flag-NMNAT2 and SARM1(-GFP) expression vectors, as expected for a largely functional NMNAT2, although, notably, survival was consistently slightly reduced compared to co-injection with wild-type Flag-NMNAT2 and SARM1(-GFP) vectors (Fig. 4D). Furthermore, we subsequently found that T94M Flag-NMNAT2 was significantly less able to protect neurites after transection in this context (no endogenous NMNAT2) than in wild-type neurons (Fig. 4A and 4E).

Together these data show that T94M NMNAT2 has a reduced capacity to maintain neurite health compared to wild-type NMNAT2 and that its protective capacity is also more sensitive to increasing temperature. This altered functionality is likely to make a major contribution to the symptoms in the affected sisters, with the hypomorphic *NMNAT2* allele likely making axons more sensitive to otherwise sub-threshold insults or stresses.

### T94M NMNAT2 is less stable than wild-type NMNAT2

Loss of stability or loss of enzymatic activity are possible alternative explanations for the reduced capacity of T94M Flag-NMNAT2 to protect cut axons in the above assays. To investigate whether stability of T94M Flag-NMNAT2 is altered we used immunoblotting to compare its rate of turnover to that of wild-type Flag-NMNAT2 at both 37°C and 39°C in transfected HEK 293T cells.

Intriguingly, we first found that T94M Flag-NMNAT2 was more sensitive than wild-type Flag-NMNAT2 to heating in sample loading buffer (data not shown). The T94M mutation thus appears to make NMNAT2 inherently less stable even in a situation where the cell degradation machinery should be largely inactive. As a consequence, we avoided sample heating in the turnover assays, but still found steady-state levels of T94M Flag-NMNAT2 (just prior to the protein synthesis block to assess turnover) to be consistently lower than wild-type Flag-NMNAT2 (Fig. 5A and 5B). This contrasts with injected SCG neurons where steady-state expression levels appeared more similar (Fig. 4C). We also found that turnover of T94M Flag-NMNAT2 was significantly accelerated relative to wild-type Flag-NMNAT2 when cells were incubated at either 37°C or 39°C after blocking protein synthesis with 10 µM emetine (Fig. 5A and 5C). Notably, turnover of both proteins was increased by heating and, as such, the decay curve of wild-type Flag-NMNAT2 at 39°C closely matches that of T94M Flag-NMNAT2 at 37°C (Fig. 5C). Intriguingly, this overlap between decay curves mirrors the similarity in the degree of protection conferred to cut neurites by wild-type Flag-NMNAT2 at 37°C and T94M Flag-NMNAT2 at 39°C (Fig. 4A).

**Figure 5.**
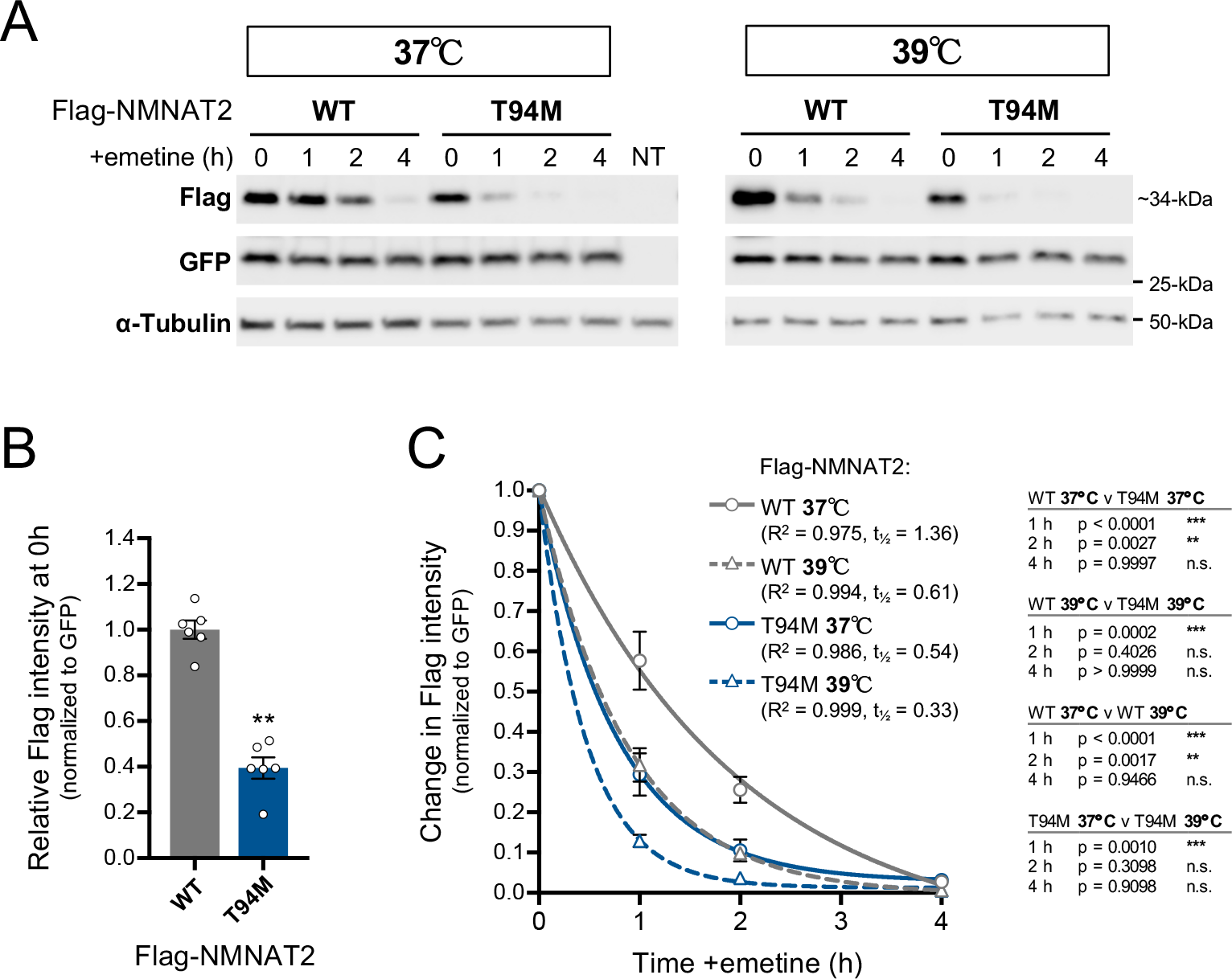
Reduced stability of T94M NMNAT2 in HEK 293T cells. (A) Representative immunoblots (of n = 3 per temperature) of extracts of HEK 293T cells co-transfected with expression vectors for wild-type (WT) or T94M Flag-NMNAT2 and eGFP at the indicated times after addition of 10 µM emetine with incubation at either 37°C or 39°C (from the time of emetine addition). Emetine was added 24 h after transfection. Non-transfected cell extract at 0h is also shown (NT). Blots were probed with Flag, eGFP and α-Tubulin antibodies. Expression of the Flag-NMNAT2 proteins was kept relatively low to avoid saturation of the protein degradation machinery by including empty vector in the transfection mix (see Materials and Methods). Co-transfected eGFP and endogenous α-Tubulin (present in transfected and non-transfected cells) are relatively stable proteins that were respectively used as references for Flag-NMNAT2 turnover (to control for transfection efficiency) and loading. (B) Relative steady-state Flag-NMNAT2 protein band intensities, normalized to eGFP, for blots described in A (0h, just before emetine addition and incubation at different temperatures). Individual values (n = 6) and means ±SEM are plotted. ** p < 0.01, Student’s *t*-test. (C) Relative turnover rates of wild-type and T94M Flag-NMNAT2 after emetine addition. Flag-NMNAT2 band intensities at each timepoint in blots described in A were normalized to co-transfected eGFP and intensities calculated as a proportion of the intensity at 0h. Means ± SEM (n = 3 per temperature) are plotted. One-phase decay curves were fitted to each data set using non-linear regression. R^2^ value and half-life (t_1/2_) are reported for each curve. Statistical comparisons are shown on the right (two-way ANOVA with Tukey’s multiple comparisons test for effects between variants).

As well as potentially making NMNAT2 inherently less stable, the T94M substitution also substantially increases the rate of turnover of the protein in cells. Accelerated loss of T94M NMNAT2 after interruption of its supply could thus contribute to its reduced ability to delay degeneration of transected neurites.

### *In vitro* characterization of human T94M NMNAT2

Given that the T94M mutant human NMNAT2 is functionally defective in cell-based axon degeneration assays, we next assessed its *in vitro* biochemical properties. Recombinant isoforms of wild type and T94M human NMNAT2 were obtained after bacterial expression and purification as His-tagged fusion proteins. The two purified enzymes (Fig. 6A) showed similar specific activity rates at 37 °C under saturating concentration of substrates, corresponding to 10.2 ± 4.8 U/mg (WT) and 13.7 ± 3.2 U/mg (T94M), in keeping with previous reports^9,12,22,23^ and were both correctly recognized by specific anti-His and anti-NMNAT2 antibodies (Fig. 6B). They were also stable after freezing at −80 °C while becoming progressively inactivated after thawing, as earlier reported^10^. However, the loss of activity was negligible in the context of our routine experiments (see Materials and Methods).

**Figure 6.**
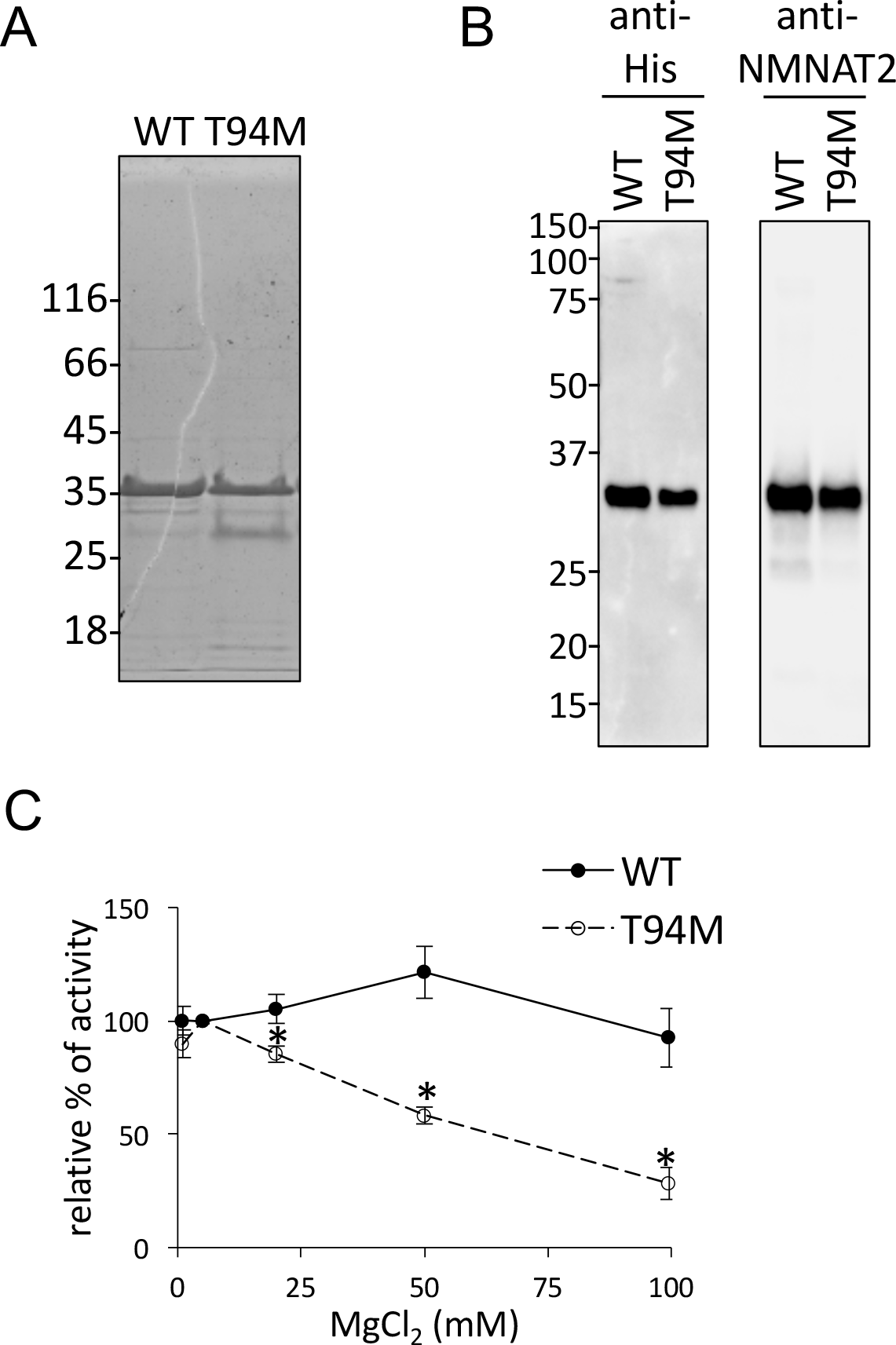
SDS-PAGE, immunoblotting and Mg^2+^-dependence of WT and T94M NMNAT2. (A) Coomassie blue stained 12 % polyacrylamide gel after running of ~2 μg each purified protein species. (B) Membrane-immobilized proteins after probing with monoclonal anti-NMNAT2 and chemiluminescence imaging. (C) Magnesium-dependent changes of rates (mean ± SEM from n = 4) referred to the value at 5 mM MgCl_2_ (arbitrary 100%). T test p values (*) for T94M *vs* WT: 0.037 at 20 mM MgCl_2_, 0.006 at 50 mM MgCl_2_, and 0.009 at 100 mM MgCl_2_.

Parallel *in vitro* assays of wild-type and T94M NMNAT2 established some differential properties arising from the mutation. First, we observed a difference in Mg^2+^-dependence. Both enzymes showed optimum activity at ~1 mM Mg^2+^, as previously reported^12^, but the mutant, unlike the wild-type, was linearly impaired at Mg^2+^ concentrations above 5 mM (Fig 6C). While such concentrations are not physiologically relevant, they are relevant to *in vitro* assays where activity is usually measured with a large excess of Mg^2+^ ions^11,12^. Thus, all subsequent assays of T94M mutant were done using a fixed MgCl_2_ concentration of 5 mM, leading to free [Mg^2+^] in low excess over [ATP].

Under these conditions, kinetic studies were carried out at 37 °C. As shown in Table 1, most kinetic parameters were similar between the two enzymes and in keeping with our previous report^12^. The main difference was a greatly increased *K*_m_ value for NMN in the T94M mutant (256.8 ± 36.7 µM) relative to wild-type (38.0 ± 6.75 µM) (p = 0.00064). Similar results were obtained using the deamidated alternative substrate NaMN in place of NMN (Supplemental Fig. 1). Thus, T94M mutant NMNAT2 has a striking ~5-fold reduced affinity towards its pyridine mononucleotide substrate which we predict would result in an increased steady-state level of NMN inside the axons of the two siblings that express only T94M NMNAT2. Supporting this, we reconstituted the main axonal pathway for NAD biosynthesis *in vitro* using NAMPT-NMNAT2 coupled reactions (Fig. 7) and evaluated in parallel the time-dependent fluctuation of Nam, NMN and NAD levels. As predicted, there was greater accumulation of NMN with T94M NMNAT2 than with the wild-type enzyme (~50 μM vs ~10 μM respectively) (Fig. 7).

**Table 1.** Kinetic parameters of human NMNAT2 WT and T94M

| Enzyme | Substrate | $K_m$<br>($\mu\text{M}$) | $K_{cat}$<br>($\text{s}^{-1}$) | $K_{cat}/K_m$<br>($\text{s}^{-1} \text{M}^{-1}$) |
| --- | --- | --- | --- | --- |
| NMNAT2 WT | ATP | $169.4 \pm 45.5$ | $8.40 \pm 2.92$ | $0.65 * 10^5$ |
| | NMN | $38.0 \pm 6.75$ | $8.40 \pm 2.92$ | $2.36 * 10^5$ |
| NMNAT2 T94M | ATP | $286.6 \pm 32.9$ | $9.20 \pm 3.80$ | $0.31 * 10^5$ |
| | NMN | $256.8 \pm 36.7 (*)$ | $9.20 \pm 3.80$ | $0.35 * 10^5$ |

Data represent Mean ± SEM of 3 independent experiments.
(\*) T test p value of 0.00064 vs the WT enzyme

**Figure 7.**
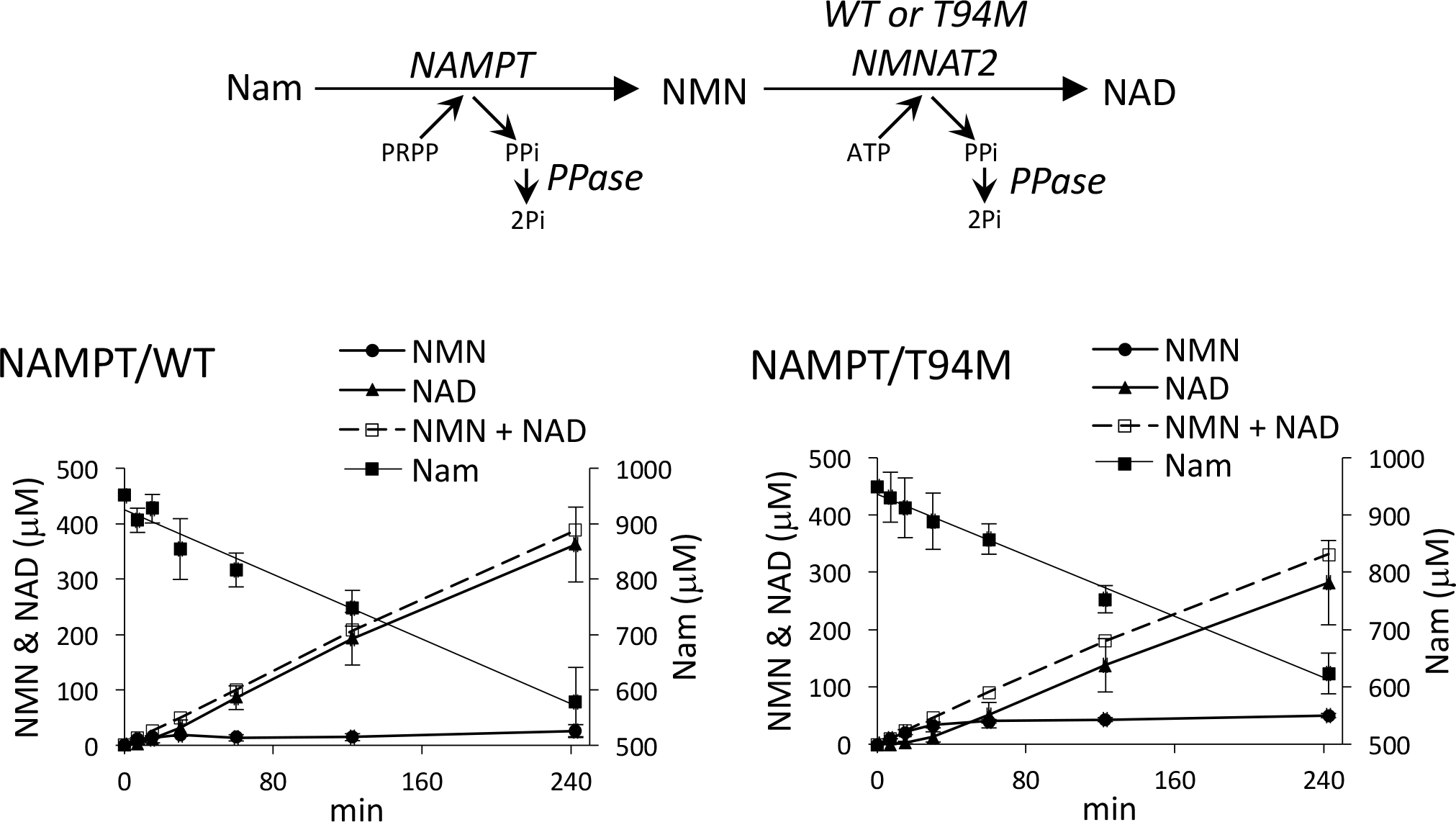
Accumulation of the NMN intermediate in NAMPT-NMNAT2 coupled reactions: comparison between WT and T94M NMNAT2. *In vitro* HPLC assays after coupling a pure recombinant murine NAMPT to either WT or T94M mutant human NMNAT2. The NAD biosynthesis starts from Nam (nicotinamide) and PPase (inorganic pyrophosphatase) was added to prevent the reverse reactions (see the top scheme). Data represent mean ± SD of 2 independent experiments. The two time-course analyses show, under similar NAD synthesis rates, a higher transient accumulation of the NMN intermediate in the T94M mutant.

The effect of temperature on the activity of the purified wild-type and T94M enzymes was also evaluated (Fig. 8). T94M NMNAT2 was slightly more prone to inactivation at a range of temperatures (Fig. 8A) and its activity had a significantly reduced half-life relative to wild type NMNAT2 at 37 °C (Fig. 8B). Notably, thermal inactivation of both enzymes was delayed in the presence of substrates but T94M NMNAT2 was still relatively less stable (Fig. 8C). Finally, we found that the optimal temperature for T94M NMNAT2 activity is approximately 5°C lower than the wild-type enzyme (Fig. 8D).

**Figure 8.**
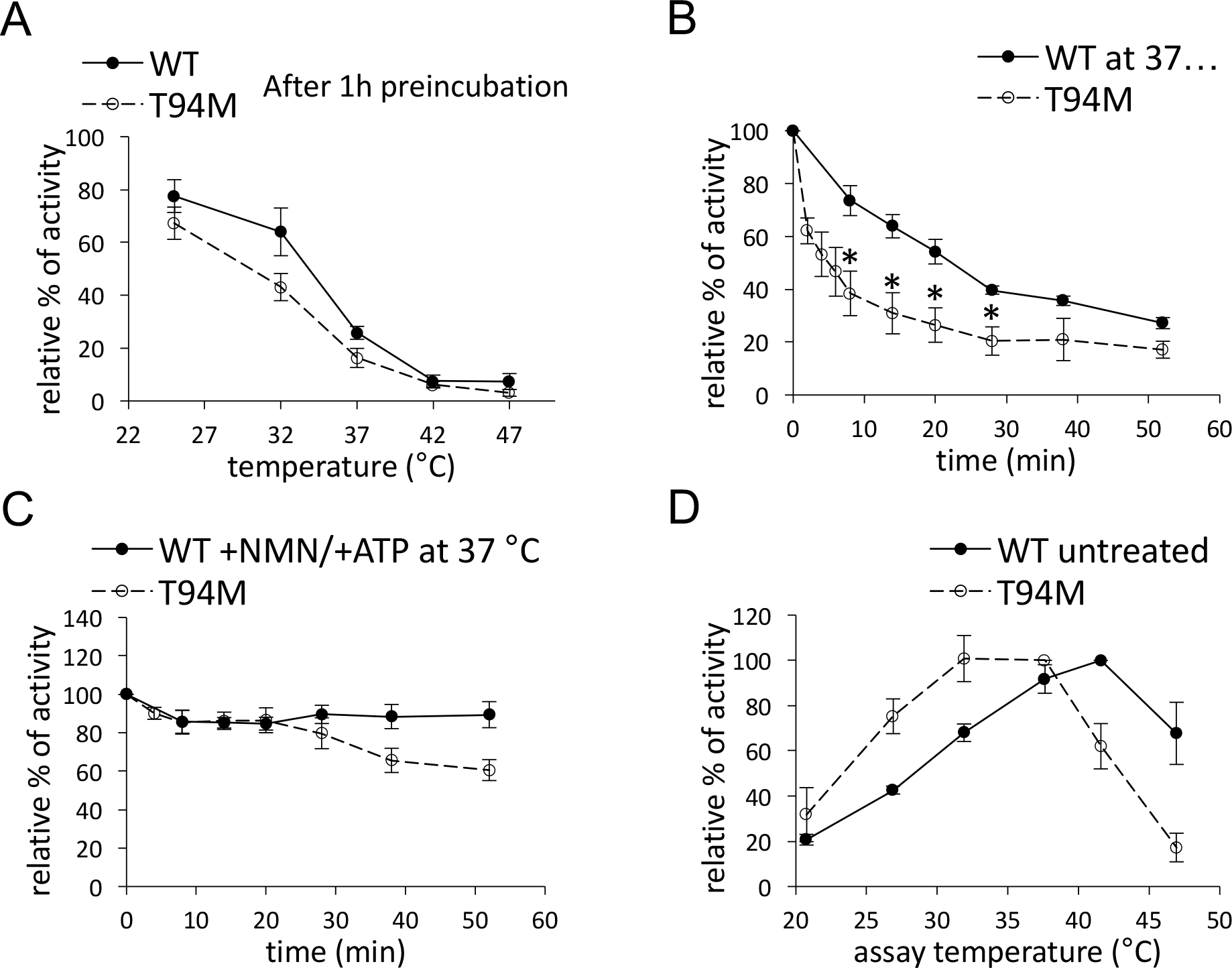
Temperature studies of WT and T94M NMNAT2. Data represent mean ± SEM of n experiments as indicated. (A) Apo-enzymes’ stability at different temperatures. Buffered enzyme solutions (40 μg/ml hNMNAT2 WT or 30 μg/ml T94M mutant in 50 mM HEPES/NaOH buffer, pH 7.5, 1 mM TCEP, 20 % glycerol) were pre-incubated for 1 hour at the indicated temperatures, then assayed at 37 °C. Values (n = 6) are referred to the untreated enzyme kept at +4 °C (arbitrary 100%). (B) Apo-enzymes’ stability at 37 °C. Enzyme solutions were incubated and collected at the indicated time intervals, then assayed at 37 °C. Values (n = 4) are referred to that of time zero (arbitrary 100%). T test p values (*) for T94M *vs* WT: 0.013 at 8 min, 0.010 at 14 min, 0.013 at 20 min, and 0.014 at 28 min. (C) Enzymes’ stability at 37 °C in the presence of substrates. Enzyme solutions supplied with 100 μM both NMN and ATP were treated and assayed as in panel B. (D) Optimal temperature. Enzyme rates were measured after warming the assay mixtures at the indicated temperatures. Values (n = 4) are referred to the relative maximal rate observed for each enzyme (arbitrary 100%). After this assay, the mixtures at 47 °C were rapidly cooled down to 37 °C then new NMNAT2 aliquots were added and rates measured again, demonstrating full recovery of the original activity. This excludes any effect caused by heating on the ancillary enzyme ADH.

Overall these data suggest that the T94M mutation has detrimental effects on both the activity and thermal stability of human NMNAT2. The predicted outcome when T94M NMNAT2 activity becomes rate-limiting in the NAD biosynthetic pathway is a rise in NMN and decline in NAD levels which, either individually, or in combination, is likely to compromise axon survival.

## DISCUSSION

In this article we describe two sisters with polyneuropathy and erythromelalgia who were found to be homozygous for an *NMNAT2* coding variant. Similar to patients who carry mutations in SCN9A, onset of symptoms was within the first decade of life and the pain attacks were accompanied by reddening and swelling of the feet and were relieved by cooling^24^. However, unlike patients with mutations in *SCN9A*, the attacks in the patients described in this article were not only provoked by heat or exercise but also by infections. Moreover, the duration of the pain attacks in the patients described here lasted from several days up to 6 weeks, while the patients with *SCN9A* mutations have attacks that last 5 minutes up to 17 hours. Finally, patients with SCN9A mutations do not have polyneuropathy, whereas it was present in both patients described here. These obvious differences in the clinical phenotype might explain why we were unsuccessful in our search for more patients with variants in *NMNAT2* among subjects with erythromelalgia as these were more typical cases of primary erythromelalgia without polyneuropathy.

Biochemical and cell-based functional assays have identified defects in T94M NMNAT2 that reinforce its candidacy as the underlying cause of the polyneuropathy with erythromelalgia in the affected siblings via a mechanism involving axon loss or dysfunction. Crucially, the metabolic consequences common to the ~5-fold increased Km value for its pyridine mononucleotide substrate and its reduced stability, both at a protein and activity level, are a predicted increase in NMN levels and a decline in NAD. This matches the situation that occurs in injured axons in vivo prior to degeneration^25^. Whilst, the altered kinetic property of T94M NMNAT2 is predicted to be stable, the intermittent nature of the phenotype in the patients likely reflects an interaction between fluctuations in body temperature and the increased thermal instability of the mutant.

NMNAT2 is to date the only mammalian isoform without a defined 3D structure but sequence alignment with other NMNATs indicates that T94 in human NMNAT2 is flanked by two conserved residues involved in pyridine nucleotide binding, corresponding to W92 and T95^26,27^. However, a T94 equivalent is not conserved in other NMNATs so its importance for substrate interaction was previously unclear. The change in substrate affinity of human T94M NMNAT2 suggests T94 is either directly involved in substrate interaction or that the amino acid substitution interferes with substrate binding to nearby residues. Altered thermal stability of the mutant protein also suggests an important role for this residue in protein folding.

All previous data on NMNAT2 mutant phenotypes come from mouse studies where expression is either reduced or ablated. The T94M mutation only partly disrupts the function of NMNAT2, so T94M homozygosity in the patients is arguably more similar to mice with heterozygous levels (~50%) or sub-heterozygous levels (~30%) of NMNAT2, which are viable into old age, than to mice homozygous for a silenced allele where a failure of nerve growth precludes survival^6,28^. Intriguingly, there are strong parallels between the patients described here and mice with sub-heterozygous levels of NMNAT2. Specifically, both have a clear, early-onset sensory phenotype involving loss of myelinated sensory axons in peripheral nerves and altered temperature sensation^28^. The absence of this phenotype in mice with heterozygous NMNAT2 levels outwardly suggests that episodes of erythromelalgia in the patients requires reduction in NMNAT2 activity to sub-heterozygous levels. However, the greater length of human axons could make them more sensitive to smaller reductions in activity than in the mice and there are indications of compensatory responses in the heterozygous mice^6^ which may be absent in patients expressing a missense variant.

The accompanying paper (Lukacs et al) reports a more severe and lethal phenotype associated with near-complete biallelic loss-of-function of NMNAT2. In common with the T94M phenotype, this is consistent with an axon loss phenotype (or failure of axon growth) affecting multiple neuron types, as also observed in both the complete null and knockdown mice^6,18,28^. The severity of the two human phenotypes correlates well with the degree of loss-of-function with the corresponding models in mice. Thus, we propose that these mutations represent different points in an allelic series and that there is a likelihood also of other human NMNAT2 phenotypes. Possible phenotypes include severe sensory and motor neuropathies intermediate between the two currently reported and subclinical phenotypes that predispose to adult-onset, acquired axonal disorders such as chemotherapy-induced peripheral neuropathy, or modifying axonal outcomes in conditions such as multiple sclerosis. A reduction in NMNAT2 expression has also been linked to tauopathy in mice^29^ and to reduced cognitive function in humans^30^. In addition, the human *SARM1* locus has been associated with sporadic ALS^31^ and SARM1 acts downstream of NMNAT2 in a Wallerian-like axon degeneration pathway. If warmth, infection or stress promote axon loss by reducing activity of the T94M variant NMNAT2 in the patients to cause the polyneuropathy with erythromelalgia phenotype, there are additional questions to address regarding the mechanism linking axon loss to pain. One hypothesis could be that an inflammatory reaction to degenerating axons somehow activates neighboring thermal nociceptors, much like partial nerve degeneration induced by chronic constriction injury causes pain through surviving receptors.

Finally, from a therapeutic viewpoint, while there is as yet no clinically available means to block axon degeneration caused by NMNAT2 deficiency, this is an area of considerable current research^17^ and genetic removal of SARM1 has already been shown to strongly rescue axons lacking NMNAT2^21,32^.

## Acknowledgements

The work was supported by grants from the German Research Foundation (Ga354/14-1), Medical Research Council grant MR/N004582/1, a Biotechnology and Biological Sciences Research Council Institute Strategic Programme Grant to the Babraham Institute Signalling ISPG and the John and Lucille van Geest Foundation.

---

**Supplemental Figure 1.**
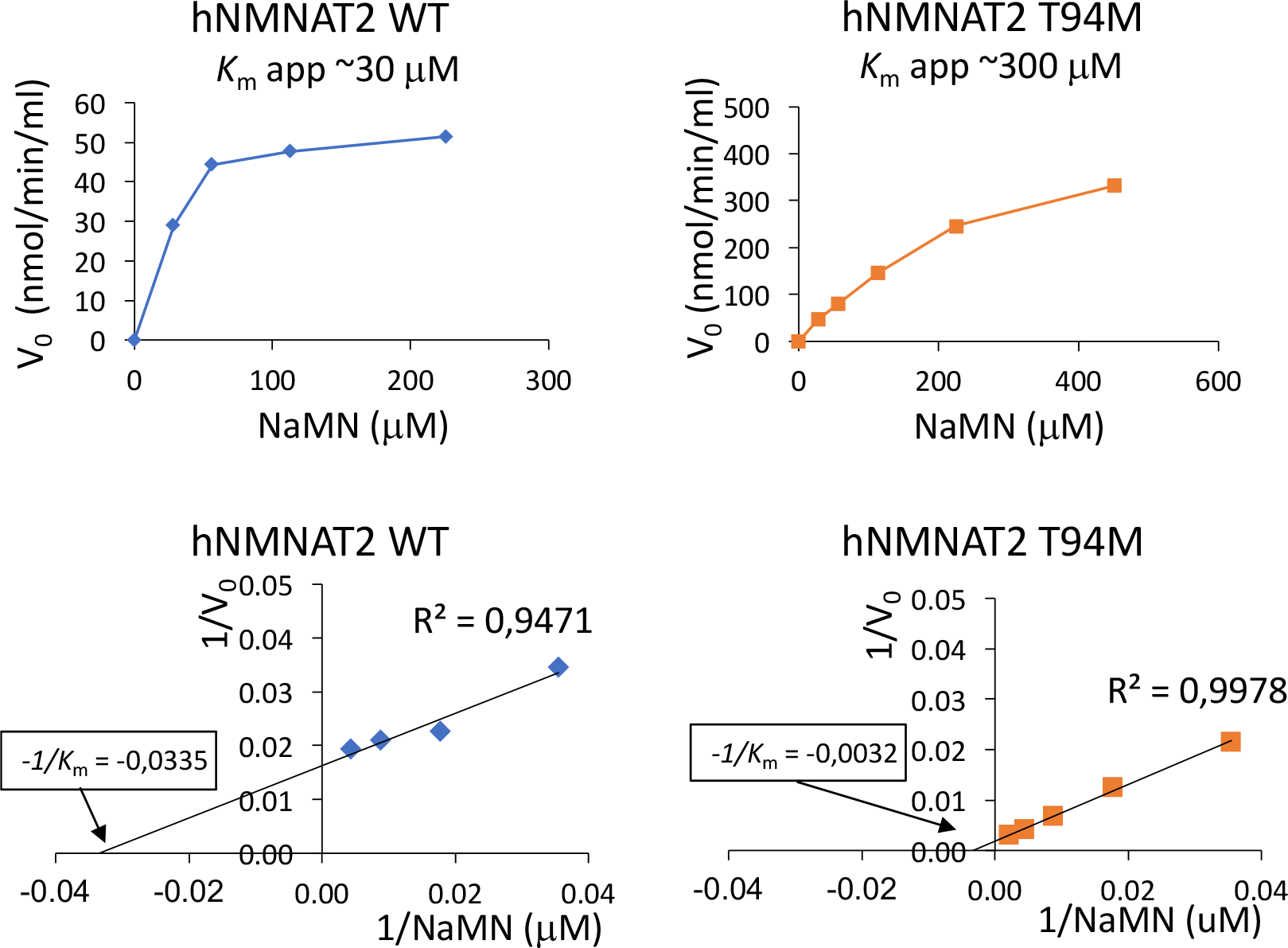
Affinity of human NMNAT2 WT and T94M for the alternative pyridine substrate NaMN. Upper panels, primary plots of initial rates measured at variable NaMN and fixed 250 μM ATP. Lower panels, the corresponding Lineweaver-Burk plots.

## REFERENCES

1. Mitchell, S.W. (1878). On a rare vaso-motor neurosis of the extremities and on the maladies with which it may be confounded. Am J Med Sci 76, 2–36.

2. Reed, K.B., and Davis, M.D. (2009). Incidence of erythromelalgia: a population-based study in Olmsted County, Minnesota. J Eur Acad Dermatol Venereol 23, 13–15.

3. Layzer, R.B. (2001). Hot feet: erythromelalgia and related disorders. J Child Neurol 16, 199–202.

4. Dib-Hajj, S.D., Rush, A.M., Cummins, T.R., Hisama, F.M., Novella, S., Tyrrell, L., Marshall, L., and Waxman, S.G. (2005). Gain-of-function mutation in Nav1.7 in familial erythromelalgia induces bursting of sensory neurons. Brain 128, 1847–1854.

5. Tang, Z., Chen, Z., Tang, B., and Jiang, H. (2015). Primary erythromelalgia: a review. Orphanet J Rare Dis 10, 127.

6. Gilley, J., Adalbert, R., Yu, G., and Coleman, M.P. (2013). Rescue of peripheral and CNS axon defects in mice lacking NMNAT2. J Neurosci 33, 13410–13424.

7. Gilley, J., and Coleman, M.P. (2010). Endogenous Nmnat2 is an essential survival factor for maintenance of healthy axons. PLoS Biol 8, e1000300.

8. Schindelin, J., Arganda-Carreras, I., Frise, E., Kaynig, V., Longair, M., Pietzsch, T., Preibisch, S., Rueden, C., Saalfeld, S., Schmid, B., et al. (2012). Fiji: an open-source platform for biological-image analysis. Nat Methods 9, 676–682.

9. Brunetti, L., Di Stefano, M., Ruggieri, S., Cimadamore, F., and Magni, G. (2010). Homology modeling and deletion mutants of human nicotinamide mononucleotide adenylyltransferase isozyme 2: new insights on structure and function relationship. Protein Sci 19, 2440–2450.

10. Orsomando, G., Cialabrini, L., Amici, A., Mazzola, F., Ruggieri, S., Conforti, L., Janeckova, L., Coleman, M.P., and Magni, G. (2012). Simultaneous single-sample determination of NMNAT isozyme activities in mouse tissues. PLoS One 7, e53271.

11. Balducci, E., Emanuelli, M., Raffaelli, N., Ruggieri, S., Amici, A., Magni, G., Orsomando, G., Polzonetti, V., and Natalini, P. (1995). Assay methods for nicotinamide mononucleotide adenylyltransferase of wide applicability. Anal Biochem 228, 64–68.

12. Sorci, L., Cimadamore, F., Scotti, S., Petrelli, R., Cappellacci, L., Franchetti, P., Orsomando, G., and Magni, G. (2007). Initial-rate kinetics of human NMN-adenylyltransferases: substrate and metal ion specificity, inhibition by products and multisubstrate analogues, and isozyme contributions to NAD+ biosynthesis. Biochemistry 46, 4912–4922.

13. Grolla, A.A., Torretta, S., Gnemmi, I., Amoruso, A., Orsomando, G., Gatti, M., Caldarelli, A., Lim, D., Penengo, L., Brunelleschi, S., et al. (2015). Nicotinamide phosphoribosyltransferase (NAMPT/PBEF/visfatin) is a tumoural cytokine released from melanoma. Pigment Cell Melanoma Res 28, 718–729.

14. Mori, V., Amici, A., Mazzola, F., Di Stefano, M., Conforti, L., Magni, G., Ruggieri, S., Raffaelli, N., and Orsomando, G. (2014). Metabolic profiling of alternative NAD biosynthetic routes in mouse tissues. PLoS One 9, e113939.

15. Lei, X.F., Fu, W., Kim-Kaneyama, J.R., Omoto, T., Miyazaki, T., Li, B., and Miyazaki, A. (2016). Hic-5 deficiency attenuates the activation of hepatic stellate cells and liver fibrosis through upregulation of Smad7 in mice. J Hepatol 64, 110–117.

16. Kim-Kaneyama, J.R., Takeda, N., Sasai, A., Miyazaki, A., Sata, M., Hirabayashi, T., Shibanuma, M., Yamada, G., and Nose, K. (2011). Hic-5 deficiency enhances mechanosensitive apoptosis and modulates vascular remodeling. J Mol Cell Cardiol 50, 77–86.

17. Conforti, L., Gilley, J., and Coleman, M.P. (2014). Wallerian degeneration: an emerging axon death pathway linking injury and disease. Nat Rev Neurosci 15, 394–409.

18. Hicks, A.N., Lorenzetti, D., Gilley, J., Lu, B., Andersson, K.E., Miligan, C., Overbeek, P.A., Oppenheim, R., and Bishop, C.E. (2012). Nicotinamide mononucleotide adenylyltransferase 2 (Nmnat2) regulates axon integrity in the mouse embryo. PLoS One 7, e47869.

19. Milde, S., Gilley, J., and Coleman, M.P. (2013). Subcellular localization determines the stability and axon protective capacity of axon survival factor Nmnat2. PLoS Biol 11, e1001539.

20. Yan, T., Feng, Y., Zheng, J., Ge, X., Zhang, Y., Wu, D., Zhao, J., and Zhai, Q. (2010). Nmnat2 delays axon degeneration in superior cervical ganglia dependent on its NAD synthesis activity. Neurochem Int 56, 101–106.

21. Gilley, J., Orsomando, G., Nascimento-Ferreira, I., and Coleman, M.P. (2015). Absence of SARM1 rescues development and survival of NMNAT2-deficient axons. Cell Rep 10, 1974–1981.

22. Berger, F., Lau, C., Dahlmann, M., and Ziegler, M. (2005). Subcellular compartmentation and differential catalytic properties of the three human nicotinamide mononucleotide adenylyltransferase isoforms. J Biol Chem 280, 36334–36341.

23. Raffaelli, N., Sorci, L., Amici, A., Emanuelli, M., Mazzola, F., and Magni, G. (2002). Identification of a novel human nicotinamide mononucleotide adenylyltransferase. Biochem Biophys Res Commun 297, 835–840.

24. McDonnell, A., Schulman, B., Ali, Z., Dib-Hajj, S.D., Brock, F., Cobain, S., Mainka, T., Vollert, J., Tarabar, S., and Waxman, S.G. (2016). Inherited erythromelalgia due to mutations in SCN9A: natural history, clinical phenotype and somatosensory profile. Brain 139, 1052–1065.

25. Di Stefano, M., Nascimento-Ferreira, I., Orsomando, G., Mori, V., Gilley, J., Brown, R., Janeckova, L., Vargas, M.E., Worrell, L.A., Loreto, A., et al. (2015). A rise in NAD precursor nicotinamide mononucleotide (NMN) after injury promotes axon degeneration. Cell Death Differ 22, 731–742.

26. Rodionova, I.A., Zuccola, H.J., Sorci, L., Aleshin, A.E., Kazanov, M.D., Ma, C.T., Sergienko, E., Rubin, E.J., Locher, C.P., and Osterman, A.L. (2015). Mycobacterial nicotinate mononucleotide adenylyltransferase: structure, mechanism, and implications for drug discovery. J Biol Chem 290, 7693–7706.

27. Zhou, T., Kurnasov, O., Tomchick, D.R., Binns, D.D., Grishin, N.V., Marquez, V.E., Osterman, A.L., and Zhang, H. (2002). Structure of human nicotinamide/nicotinic acid mononucleotide adenylyltransferase. Basis for the dual substrate specificity and activation of the oncolytic agent tiazofurin. J Biol Chem 277, 13148–13154.

28. Gilley, J., Mayer, P.R., Yu, G., and Coleman, M.P. (2019). Low levels of NMNAT2 compromise axon development and survival. Hum Mol Genet 28, 448–458.

29. Ljungberg, M.C., Ali, Y.O., Zhu, J., Wu, C.S., Oka, K., Zhai, R.G., and Lu, H.C. (2012). CREB-activity and nmnat2 transcription are down-regulated prior to neurodegeneration, while NMNAT2 over-expression is neuroprotective, in a mouse model of human tauopathy. Hum Mol Genet 21, 251–267.

30. Ali, Y.O., Allen, H.M., Yu, L., Li-Kroeger, D., Bakhshizadehmahmoudi, D., Hatcher, A., McCabe, C., Xu, J., Bjorklund, N., Taglialatela, G., et al. (2016). NMNAT2:HSP90 Complex Mediates Proteostasis in Proteinopathies. PLoS Biol 14, e1002472.

31. Fogh, I., Ratti, A., Gellera, C., Lin, K., Tiloca, C., Moskvina, V., Corrado, L., Soraru, G., Cereda, C., Corti, S., et al. (2014). A genome-wide association meta-analysis identifies a novel locus at 17q11.2 associated with sporadic amyotrophic lateral sclerosis. Hum Mol Genet 23, 2220–2231.

32. Gilley, J., Ribchester, R.R., and Coleman, M.P. (2017). Sarm1 Deletion, but Not Wld(S), Confers Lifelong Rescue in a Mouse Model of Severe Axonopathy. Cell Rep 21, 10–16.

